# Validation of biosignatures confirms the informative nature of fossil organic Raman spectra

**DOI:** 10.1101/2021.02.07.430162

**Authors:** Jasmina Wiemann, Derek E. G. Briggs

## Abstract

Raman spectroscopy has facilitated rapid progress in the understanding of patterns and processes associated with biomolecule fossilization and revealed the preservation of biological and geological signatures in fossil organic matter. Nonetheless six large-scale statistical studies of Raman spectra of carbonaceous fossils, selected from a number of independent assessments producing similar trends, have been disputed. Alleon et al. (*21*) applied a wavelet transform analysis in an unconventional way to identify frequency components contributing to two baselined spectra selected from these studies and claimed similarities with a downloaded edge filter transmission spectrum. On the basis of indirect comparisons and qualitative observations they argued that all spectral features detected, including significant mineral peaks, can be equated to edge filter ripples and are therefore artefactual. Alleon et al. (*21*) extrapolated this conclusion to dispute not only the validity of n>200 spectra in the studies in question, but also the utility of Raman spectroscopy, a well established method, for analysing organic materials in general. Here we test the claims by Alleon et al. (*21*) using direct spectral comparisons and statistical analyses. We present multiple independent lines of evidence that demonstrate the original, biologically and geologically informative nature of the Raman spectra in question. We demonstrate that the methodological approach introduced by Alleon et al. (*21*) is unsuitable for assessing the quality of spectra and identifying noise within them. Statistical analyses of large Raman spectral data sets provide a powerful tool in the search for compositional patterns in biomaterials and yield invaluable insights into the history of life.

## 1. Introduction

Raman spectroscopy is an established method for characterizing the total chemical composition of potentially heterogenous materials (*1, 2*). Applications of non-destructive, *in situ* Raman microspectroscopy have resulted in rapid progress in understanding biomolecule fossilization and the detection of biosignatures based on comparative statistical analyses of fossil organic matter (*3-15*). The *in situ* approach facilitates rapid non-destructive analysis of surface-cleaned bio- and geomaterials without requiring time-consuming extractions of the organic matter. Raman spectroscopy not only characterizes molecular functional groups (small molecular units with distinct chemical properties) in highly heterogenous fossil organic matter, but also provides insights into higher-level structural organization by detecting intermolecular and bioinorganic (i.e., organo-mineral) interactions (*1, 2*). Raman scattering can be induced by excitation wavelengths ranging from the ultraviolet, to visible (such as 532 nm) and the near-infrared range, and all of these approaches produce reliable insights into patterns in the composition of biomaterials (quantitatively compared in *16*). Raman spectra obtained under identical conditions for a given sample set can be standard-baselined, normalized, and analyzed by means of multivariate statistics (*16-20*). Ordination methods are most commonly used for comparative spectral analyses in the search for patterns in the total organic composition of complex biological samples (*16-20*). A number of advantages, including time-efficient non-destructive analysis, wide availability and low operating cost of equipment, and utility of the results, make Raman and complementary types of light spectroscopy ideal for initial molecular assessments of paleontological and geological materials (*3-15*).

A diversity of studies conducted by different laboratories (*3-15*) have recovered very similar patterns in the molecular makeup of fossil organic matter in independently acquired *in situ* Raman spectra. However, a recent Perspective by Alleon et al. (*21*) claimed that the results of a selected subset of these studies (*3, 4, 6, 11, 12, 15*) are invalid, based on the assertion that published spectra, despite resembling previous independent Raman analyses (*7-10, 13, 14*), are the result of instrumental artefacts reflecting sinusoidal edge filter ripples. The disputed studies are based on *in situ* Raman spectra for a diversity of modern, experimentally matured, and fossil tissues of different types, as well as sedimentary host rocks. The spectra were obtained under identical conditions within each individual project. Patterns in the total organic composition of samples were revealed by multivariate statistical analyses (primarily by ordination methods) of relative intensities at Raman shifts corresponding to relevant organic functional groups (*3, 4, 6, 11, 12, 15*). The statistical analyses revealed distinct clusters corresponding to fossil and sediment spectra, and showed that the relative positions of modern, experimentally matured, and fossil data align in a manner reflecting increased organic alteration (*3, 4, 6, 11, 12, 15*). Even when spectra from different localities, tissue types, and metamorphic regimes are treated together, biologically and geologically meaningful clustering is clearly evident (*3, 4, 6, 11, 12, 15*).

## 2. The supposed appearance of non-informative artefacts in modern and fossil whole-tissue Raman spectra

Alleon et al. (*21*) presented different observations and arguments in support of their assertion that the disputed spectra are artefactual in nature. Their arguments relied primarily on a frequency contribution analysis which, as far as we can determine, has not been applied by others to baselined Raman spectra. They based their conclusions on just two (< 1% of the disputed data set) of more than 200 spectra analyzed in the publications they claim are invalid (*3, 4, 6, 11, 12, 15*). Their key observations and inferences can be summarized as follows.

### 2.1. An edge filter transmission spectrum shows major contributions of 64 cm^-1^ and 128 cm^-1^ frequency contributions

Alleon et al. (*21*) downloaded a freely available RazorEdge filter transmission spectrum (*21*, fig. 2a), which resembles a periodic, sinusoidal wave function, from the commercial website ‘Semrock’ (www.semrock.com/FilterDetails.aspx?id=LP03-532RE-25). It is not clear why this specific transmission filter was chosen. They mapped out the contributions of various frequency components over a wide range (600-6000 cm^-1^) of the filter spectrum using a wavelet transform analysis, and found a significant contribution at 64 cm^-1^ and 128 cm^-1^ (*21*, fig. 2b, continuous red band). Periodic wavelet transform analysis (*21*), which is commonly applied to acoustic or precipitation data (reviewed in *22*), is not usually applied to Raman spectra beyond determining the ideal baselines for raw data (*23*), and is inferior to Principal Component Analyses of spectral data for assessing Raman spectral signatures (*16, 24*). We are unaware of examples in the literature where wavelet transform analysis has been applied to spectra that were already baselined and normalized, which is the nature of the published spectra analyzed by Alleon et al. (*21*). This analysis decomposes a given wave form, whether periodic or not, into its frequency components, and displays their contributions to the source wave form by quantifying the extent to which a sinusoidal peak in each frequency contribution needs to be scaled to resemble the original shape of the curve. Alleon et al. (*21*) encoded this information in the form of a heat map (figs. 1a, c, 2b, 3f). Similar frequency contribution maps were calculated for two baselined and normalized spectra the authors claimed to be invalid, those of a *Rhea* eggshell and a Carboniferous shrimp *Acanthotelson* from the Mazon Creek Lagerstätte. These spectra are part of the results of different projects: The first compared fossil and modern eggshell and used fixed baselining parameters (adaptive, 30%) optimized for more heterogenous fossils (*4*), the second compared invertebrate and vertebrate fossil spectra from a single Lagerstätte using the same baselining parameters (*12*). Alleon et al. (*21*) selected the *Rhea* eggshell as a pigmented example (*21*, fig. 1a), even though it is barely pigmented compared to the *Dromaius* eggshell analysed in the same study (*4*, fig. 1). In the case of *Acanthotelson* (*21*, fig. 1c), they cropped off the spectral area between 500-600 cm^-1^ which contains distinct, narrow organosulfur bands and sulfide mineral peaks. The frequency contribution maps (*21*, figs. 1a, c, 2b) are differently scaled on their y-axes, cover different Raman shift ranges (x-axes), and are colored based on different power scaling. Alleon et al. (*21*) claimed that patterns in the frequency contributions of the two disputed spectra resemble those of the edge filter, signatures that would be considered artefacts, without providing direct comparisons. They observed only patchy contributions of various frequency components (*21*, figs. 1a, c, red colors) in different regions of the disputed spectra, in contrast to the narrow red band in the edge filter spectrum. Major contributions of the 64 cm^-1^ and 128 cm^-1^ frequency components differ substantially between the disputed *Rhea* and *Acanthotelson* spectra although Alleon et al. considered them to be similar (*21*). The *Rhea* map reveals a range of patchy low and high frequency contributions, whereas the 64-128 cm^-1^ frequency contributions in the *Acanthotelson* spectrum (*21*, fig. 1c, dark red areas) are only present at the positions associated with broad -C-C-/-C=C-/-C-N-(1350 – 1650 cm^-1^) Raman bands, which are commonly detected in carbonaceous fossils containing unordered organic carbon (*3-15, 26-33*).

**Fig 1:**
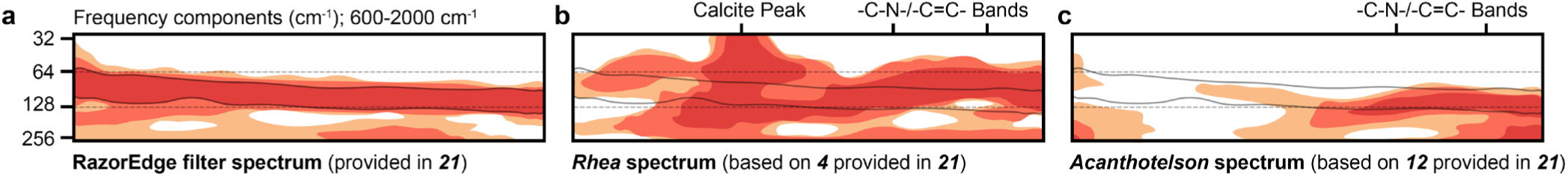
Plot of equally scaled (on x- and y-axis) **a**. RazorEdge filter (‘Semrock’), **b**. *Rhea*, **c**. *Acanthotelson* spectra frequency component maps between 600-2000 cm^-1^ Raman shift. Maps show the major contributions (*21*, figs. 1-3, red-orange colors). Peaks and bands with particularly high relative intensities are labeled.

**Fig 2:**
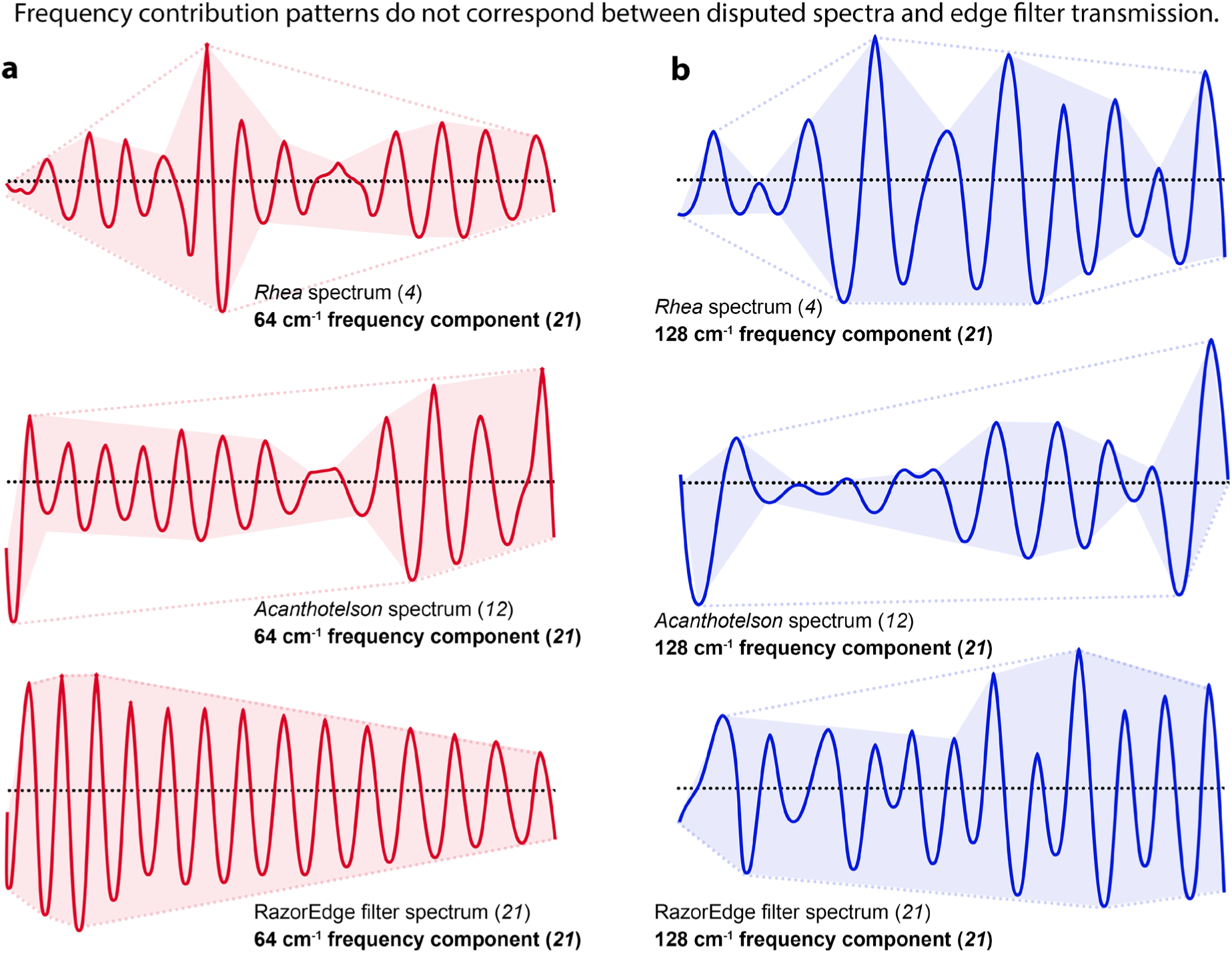
Plot of equally scaled **a**. 64 cm^-1^ and **b**. 128 cm^-1^ frequency components of the *Rhea* eggshell and Mazon Creek *Acanthotelson* spectra, and the RazorEdge filter transmission spectrum (‘Semrock’, *21*) between 600-2000 cm^-1^ Raman shift. The red and blue shaded areas highlight trends in the energy distribution of frequency contributions; dotted outlines represent convex hulls.

### 2.2. Summed frequency components closely resemble their original source spectrum

Prompted by the major contribution of almost exclusively 64 and 128 cm^-1^ frequency components to the edge filter spectrum and the suggested similarity to frequency contribution maps of the two disputed spectra, Alleon et al. (*21*) subsampled the edge filter spectrum, the two disputed spectra, and their newly analyzed spectrum of the Mazon Creek shrimp *Peachocaris* (*21*, fig. 1a) into their 64 cm^-1^ (*21*, fig. 2c, red line) and 128 cm^-1^ (*21*, fig. 2c, blue line) frequency components. They superimposed the subsampled frequency functions for a subset of their own spectra, and plotted the summed curves (lowest line in *21*, figs. 2c, 4) which closely resemble the source spectra (*21*, figs. 2b, 3d, e). It should be noted that even though the extracted 64 cm^-1^ and 128 cm^-1^ frequency contributions (*21*, fig. 1b, d, fig. 2c, fig. 4) are identically colored, they are unrelated. The blue and red frequency contributions (*21*, figs. 1b, d) are calculated directly from the disputed spectra, and do not represent the frequency functions calculated from the edge filter spectrum (*21*, fig. 2b).

The resemblance of the summed 64 cm^-1^ and 128 cm^-1^ frequency contributions to the original filter transmission spectrum led Alleon et al. (*21*) to suggest that the disputed spectra must likewise be composed of edge filter signatures, i.e., artefacts. However, the sum of two substantially distinct frequency components obtained through decomposition of a spectrum (or wave form) will inevitably resemble the original spectrum. Any combination of two distinct spectral frequency components with a wavelength smaller than that of the broadest band in the Raman spectrum (independently established ‘G-band’ in highly degraded fossil organic matter, -C=C-/-C-N-bands in less degraded fossil organic matter at 1603 cm^-1^: up to 250 cm^-1^ base width) would closely resemble the original spectrum upon re-composition (see *23* for an illustration of the decomposition process).

Alleon et al. (*21*) did not compare the 64 cm^-1^ and 128 cm^-1^ frequency contributions of the edge filter spectrum directly with those calculated from the two disputed Raman spectra. They suggested, however, that patterns in the extracted frequency contributions of these two spectra resemble patterns in the edge filter spectrum. Alleon et al. suggested that 64 cm^-1^ frequencies contribute at wavenumbers below 1000 cm^-1^, whereas 128 cm^-1^ frequency contributions are dominant at wavenumbers above 1000 cm^-1^ (*21*). A direct assessment of this claim is complicated by the different Raman shifts plotted for the frequency components extracted from the edge filter (600-6000 cm^-1^), *Rhea* (200-3000 cm^-1^) and *Acanthotelson* (600-2000 cm^-1^) spectra (*21*). When Alleon et al. (*21*) subtracted the summed 64 cm^-1^ and 128 cm^-1^ frequency components from the original spectra (*21*, figs. 1b, d) virtually no signal remained, not even the significant D- and G-bands (-C=C- and -C-N-functional groups in unmetamorphosed carbonaceous residues) associated with unordered organic matter (*3-15, 25-32*). Alleon et al. (*21*) did not discuss this unlikely result.

Based on the supposed similarity between the frequency contribution maps of the edge filter spectrum and the two disputed spectra, and their observation that the summed frequency components closely resemble their original source spectra, Alleon et al. (*21*) concluded that all Raman spectral bands obtained from fossil, experimentally matured, and extant tissue samples, even when identical to reference spectra, independently reproduced, diverse through time, statistically distinct, or biologically informative, are edge filter artefacts. Although this claim impugns the results of numerous investigations employing Raman spectroscopy to investigate biomaterials (e.g., *3-15, 26-33*), Alleon et al. (*21*) focused exclusively on the work of Wiemann and colleagues (*3, 4, 6, 11, 12, 15*) ignoring similar spectra published in other studies (*5, 7-10, 13-14, 30-31*).

### 2.3. New spectra of comparable fossils analysed by Alleon et al. (*21*) do not yield significant organic signals

Alleon et al. (*21*) presented a set of five raw spectra (*21*, fig. 3d) obtained from two fossils (*Peachocaris*, Carboniferous, Mazon Creek; *Cretapenaeus*, Cretaceous, Morocco), their sediment matrices, and a dried modern shrimp of the species *Neocaridina* (*21*, fig. 3a-c). The samples from the fossil specimens (*21*, fig. 3a,b, circles) show no sign of coinciding with carbonaceous material which is necessary to obtain an organic signal (*12*). The steep baseline leading to higher wavenumbers and the lack of any intense, sharp mineral or organic peaks (the mineral peaks are uncharacteristically weak: compare *24*, or *3, 4, 6, 11, 12, 15*) indicate that these spectra are dominated by background fluorescence (*21*, fig. 3d). Notably, the dried shrimp did not yield any signals related to chitin in the cuticle (see e.g., *34*). These spectra were then baselined, but not normalized (*21*, fig. 3e). By subjecting their own spectra to adaptive baselining, Alleon et al. (*21*) emphasized how exaggerated adaptive baselining amplifies edge filter artefacts – it is not clear, however, how baselining could affect interpretations based on the statistical evaluation of identically treated spectra, as in the disputed studies. Alleon et al. do not provide any assessments that could corroborate their claim. The spectrum obtained from the Mazon Creek fossil *Peachocaris* was subjected to a wavelet transform analysis (*21*, fig. 3f), as were the edge filter spectrum (*21*, fig. 2b), and the disputed spectra (*21*, fig. 1a,c). Alleon et al. (*21*) decomposed the spectrum into its 64 cm^-1^ and 128 cm^-1^ frequency components which, when summed, yielded a spectrum resembling the original (see Section 2.1). Alleon et al. (*21*) did not plot their own spectra against the published spectra that they claimed are invalid, relying on similarities to make their case.

**Fig 3:**
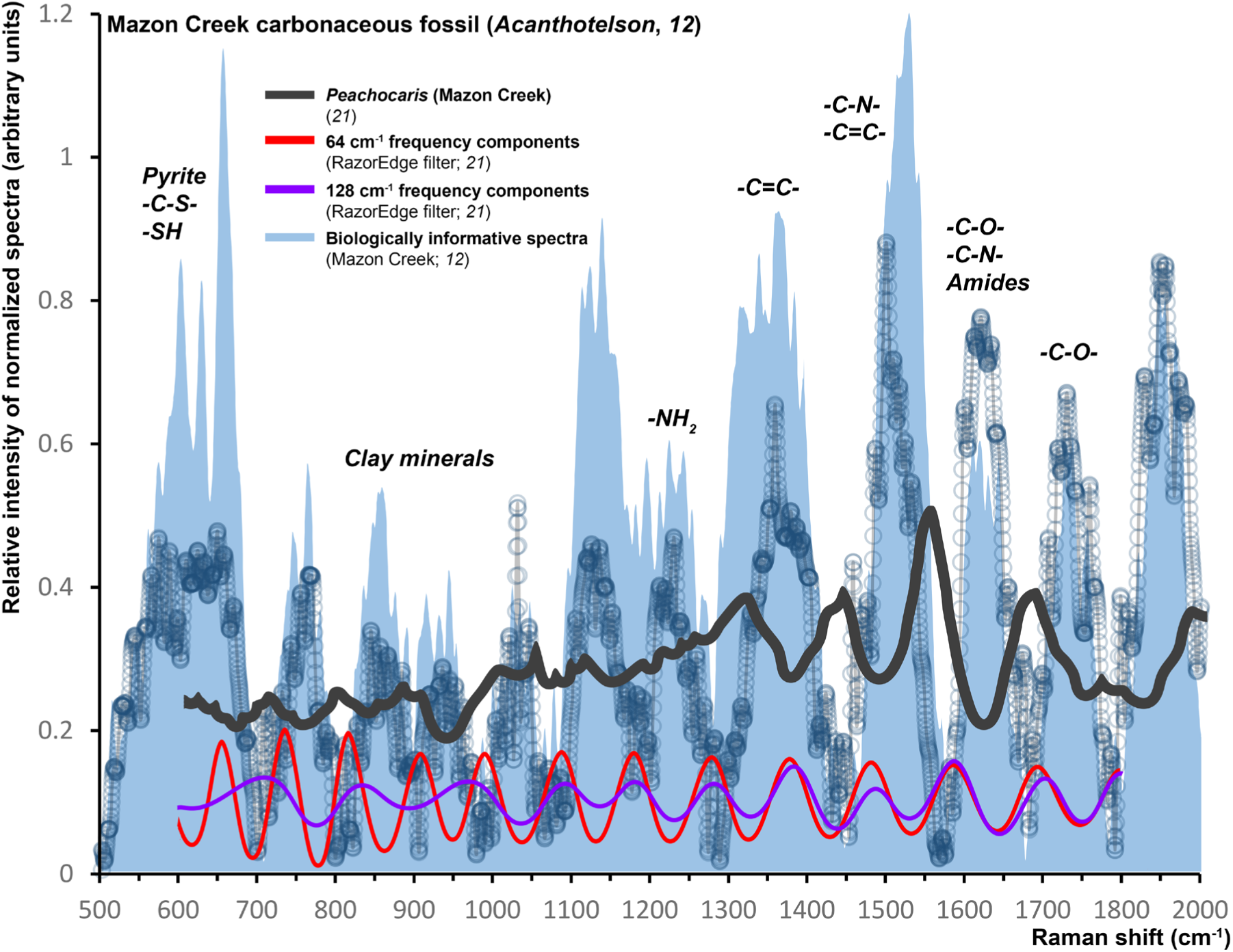
Comparison of Raman spectra of our Mazon Creek carbonaceous fossils (*12*), the Mazon Creek *Peachocaris* analyzed and interpreted by Alleon et al. (*21*) as artefactual, and the ‘Semrock’ sinusoidal edge filter 64 cm^-1^ and 128 cm^-1^ frequencies over the 600-2000 cm^-1^ range. The blue shaded area represents the diversity of our Mazon Creek fossil spectra (*12*) all of which were obtained with 10 technical replicates, normalized, and subjected to adaptive baselining (30%, SpectraGryph).

**Fig 4:**
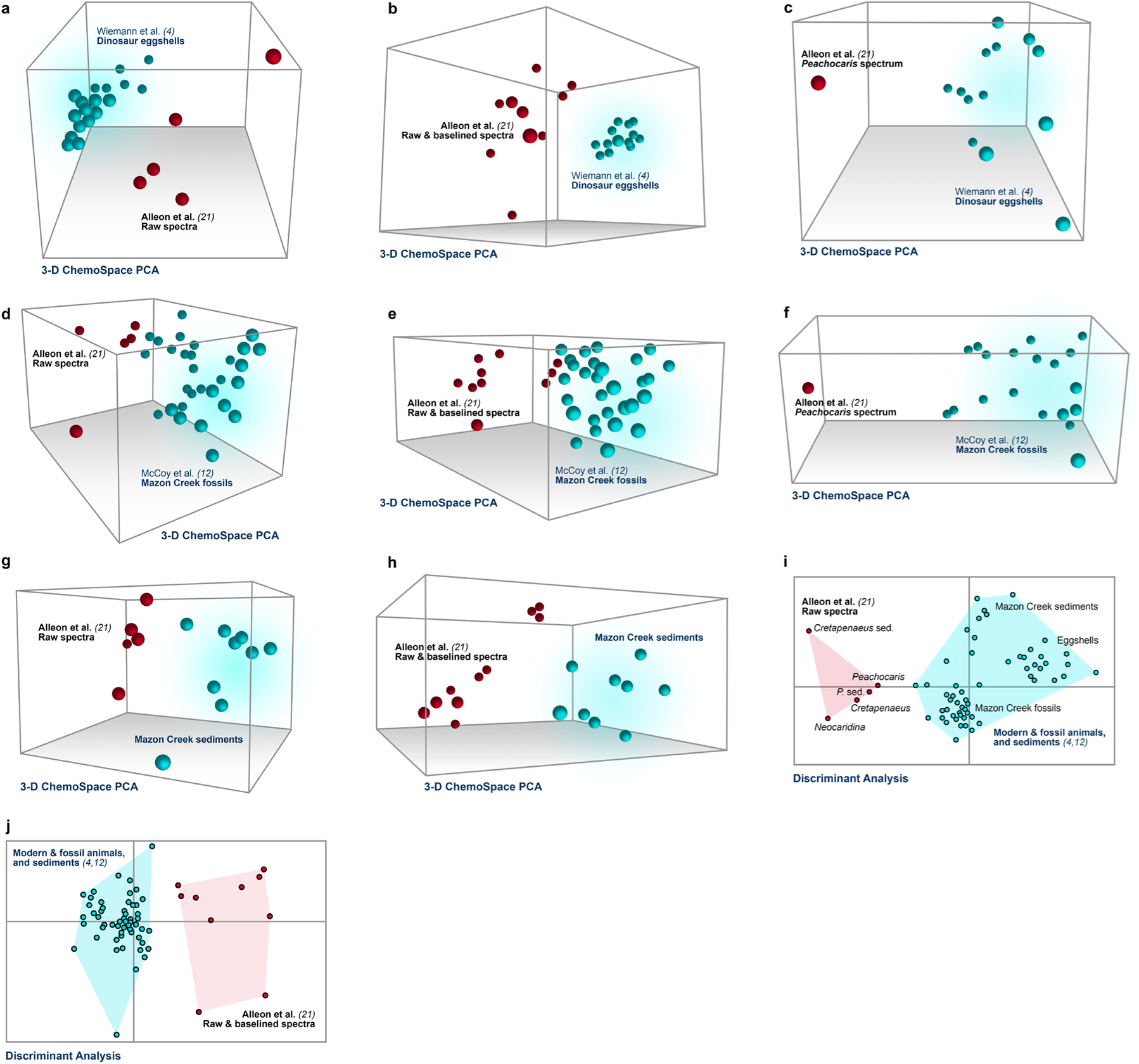
ChemoSpace Principal Component Analyses (PCA) and Discriminant Analyses of raw and baselined spectra from Alleon et al. (red data points; *21*, available on Dryad), and spectra associated with the disputed studies (blue data points; *4, 12*). Spectra were baselined and normalized together, before relative intensities were selected at n=26 band positions (n=24 band positions published in Wiemann et al. 2020 (*11*) and phosphate and carbonate bands). The 3D ChemoSpaces are plotted from scores across PC 1-3. All spectra were baselined and normalized together for each ChemoSpace PCA, and Discriminant Analysis. **a**. ChemoSpace PCA of fossil and modern eggshell spectra (n=18) (*4*) and raw spectra (n=5) from *21*. **b**. ChemoSpace PCA of fossil dinosaur eggs (n=13) (*4*) and raw (n=5) and baselined spectra (n=5) from *21*. **c**. ChemoSpace PCA of fossil dinosaur eggs (n=13) (*4*) and the *Peachocaris* spectrum interpreted as artefactual (*21*). **d**. ChemoSpace PCA of Mazon Creek vertebrate and invertebrate fossils (n=31) (*12*) and raw spectra (n=5) from *21*. **e** ChemoSpace PCA Mazon Creek vertebrate and invertebrate fossils (n=31) (*12*) and raw (n=5) and baselined spectra (n=5) from *21*. **f**. ChemoSpace PCA of Mazon Creek vertebrate and invertebrate fossils (n=31) (*12*) and the *Peachocaris* spectrum interpreted as artefactual (*21*). **g**. ChemoSpace PCA of new Mazon Creek sediment spectra (n=9) (associated with fossils in *12*) and raw spectra (n=5) from *21*. **h**. ChemoSpace PCA of new Mazon Creek sediment spectra (n=9) (associated with fossils in *12*) and raw (n=5) and baselined spectra (n=5) from *21*. **i**. Discriminant Analysis of all spectra (n=31, n=9, n=18) from the disputed studies (*4, 12*) and and raw spectra (n=5) from *21*. **j**. Discriminant Analysis of all spectra (n=31, n=9, n=18) from the disputed studies (*4, 12*) and raw (n=5) and baselined spectra (n=5) from *21*. Abbreviations: sed. = Sediment; P. sed. = *Peachocaris* sediment.

Alleon et al. (*21*) argued that the poor signal-to-noise ratio in their own spectra (*21*, fig. 3) corroborates their concerns about the spectra they dispute (*3, 4, 6, 11, 12, 15*) even though their new spectra do not resemble those of carbonaceous fossils published by a number of different laboratories (*3-15, 26-29, 32-33*). Nor did they apply statistical methods to their data or discuss the results of other studies that included analyses of 40-100 sample spectra which clearly show consistent diagnostic differences between different samples (*3, 4, 6, 11, 12, 15*, but also *5, 9*).

## 3. Raman spectra are a powerful source of chemical, biological and geological information stored in fossil organic matter, particularly when combined with quantitative comparisons

The claim by Alleon et al. (*21*) that more than 200 spectra published in a selected subset of studies (*3, 4, 6, 11, 12, 15*) represent artefacts resulting from edge filter ripples (*21*) was largely based on just two spectra. In order to test their assertion, we considered evidence from a variety of sources.

### 3.1. Published Raman spectra do not resemble the ‘Semrock’ filter transmission spectrum

Alleon et al. (*21*) argued that the frequency contribution maps of the disputed spectra resemble that of the RazorEdge filter in showing a distinct 64-128 cm^-1^ frequency contribution (*21*, fig. 2b, continuous red band). However, their maps cover different Raman shift ranges (x-axes), and are scaled differently on the y-axis (compare *21*, figs. 1a, c to fig. 2b and fig. 3f), making comparisons complicated. Moreover, the coloration of the heat map (*21*, figs. 1a, c, 2b, 3f) is scaled on substantial differences between the disputed spectra and the RazorEdge filter transmission spectrum (compare *21*, figs. 1a, c to fig. 2b), whereas no power scale is provided for the heat map of their *Peachocaris* spectrum (*21*, fig. 3). Alleon et al. (*21*) presented no direct comparison of their ‘Semrock’ filter transmission spectrum, their own spectra, and the disputed spectra, and differences in the scales on the plots of Raman shifts (compare *21*, figs. 1a, c, 2, and 3f) make comparisons difficult.

Alleon et al. (*21*) further argued that patterns in the 64 cm^-1^ and 128 cm^-1^ frequency contributions of the ‘Semrock’ RazorEdge filter are similar to patterns in those calculated from the published *Rhea* eggshell and *Acanthotelson* spectra (*21*). Alleon et al. (*21*, fig. 1d) limited their comparisons to the decomposition of our baselined spectrum into 64 cm^-1^ and 128 cm^-1^ frequency components which, when summed, predictably resemble the source spectrum. The difference in spectral ranges analyzed in Alleon et al. (*21*), and the incomplete x-axis labels provided, render any direct comparison of the presented frequency contribution data (Rhea eggshell: 500-3000 cm^-1^; *Acanthotelson* fossil: 600-2000 cm^-1^; RazorEdge filter: 600-6000 cm^-1^) inaccessible to the reader.

Here, we scaled the frequency contribution maps of the ‘Semrock’ RazorEdge spectrum and the disputed *Rhea* and *Acanthotelson* spectra in the same way, and reduced the information to the elevated contributions of the frequencies relevant to the concerns of Alleon et al. (*21*, figs. 1a, c, 2b and 3f, orange-red colors). This direct comparison of frequency contribution maps (Fig. 1) shows that the continuous (red) band of almost exclusively 64-128 cm^-1^ frequency contributions in the ‘Semrock’ RazorEdge filter spectrum (*21*, fig. 2b) is not present in either of the disputed spectra, in contrast to the claim by Alleon et al. (*21*).

Various lower and higher frequency contributions are present in the *Rhea* eggshell, whereas elevated contributions of 64-128 cm^-1^ frequency components in the fossil *Acanthotelson* spectrum are only detected in the regions of the established, broad -C-N-/-C=C-(compare D- and G-) bands. We also compare the frequency components calculated from our *Rhea* eggshell, Mazon Creek *Acanthotelson*, and the ‘Semrock’ edge filter spectra (Fig. 2) to assess the claim that patterns in the frequency contributions of our spectra resemble those of the edge filter spectrum (*21*). To aid the direct comparison of frequency contribution patterns, we shaded the general outline of the frequency contribution spectra and added convex hulls. The shaded outline allows the immediate comparison of small-scale variations in the contribution patterns, and the convex hulls highlight large-scale trends in the frequency contributions. The 64 cm^-1^ spectral frequency components of our spectra (Fig. 2a) do not mirror the constant drop in energy observed for the edge filter towards higher wavenumbers (here plotted: 600-2000 cm^-1^), and show little resemblance when compared to each other or to the 64 cm^-1^ filter frequency contributions. Alleon et al. (*21*) pointed out that the energy of 128 cm^-1^ frequency components associated with the edge filter increase towards higher wavenumbers. Patterns in the corresponding 128 cm^-1^ frequency components calculated from our spectra, however, do not align one with the other, nor with the edge filter frequency components (Fig. 2b). We plotted (Fig. 3) our Mazon Creek fossil invertebrate and vertebrate spectra (*12*) against the spectrum of *Peachocaris strongi* from Alleon et al. which they interpreted as artefactual (*21*, fig. 3f).

The lower limit for the x-axis Raman shift is not labelled in Alleon et al. (*21*, fig. 3f), so we extracted the spectral curve function between the defined 600-2000 cm^-1^ Raman shift (points which are labeled) and superimposed it on our Mazon Creek spectra (*12*).We likewise superimposed the 64 cm^-1^ and 128 cm^-1^ frequency components from the filter transmission spectrum downloaded from ‘Semrock’ – it is clear that they do not align (Fig. 3).

A comparison of these spectra (Fig. 3) shows that the Mazon Creek shrimp *Peachocaris* (*21*) resembles the sinusoidal edge filter spectrum in the rather smooth, wavy band shapes and, most prominently, in the lack of any actual peaks. None of the bands in the Mazon Creek *Peachocaris* spectrum align with those in our Mazon Creek spectra. Nor do the edge filter 64 cm^-1^ and 128 cm^-1^ frequency components align with any band detected in our Mazon Creek spectra. In addition to the lack of alignment, there is a substantial difference in the number of sharp, organic and mineral peaks contributing to the broader bands observed. The spectrum of *Peachocaris* (*21*, fig. 3), in contrast to our Mazon Creek spectra (Fig. 3), does not show any sharp organic bands combining into broad bands characteristic of heterogeneous whole-tissue samples (*3-15, 18, 26-33, 34-37*).

Alleon et al. (*21*, figs. 1-3) presented the 64 cm^-1^ and 128 cm^-1^ frequency components obtained through the decomposition of baselined spectra and the 64 cm^-1^ and 128 cm^-1^ frequency components of the ‘Semrock’ edge filter in the same colors even though they are unrelated and obtained from different sources: the frequency components shown next to the spectra (*21*, figs. 1, 3) do not represent the frequency components of the edge filter, but those calculated directly from the two disputed Raman spectra (*21*, fig. 2). Alleon et al. did not present extracted and baselined 64 cm^-1^ and 128 cm^-1^ frequency components for their *Peachocaris* spectrum (*21*, fig. 3f).

### 3.2. Wavelet transform analyses and maps are not a reliable method for assessing the quality of spectra: Significant mineral peaks and D- and G-bands are eliminated as noise

Locally elevated contributions of high frequency periodicities are the norm for independently confirmed graphite-like D- and G-bands, which are broad as a result of overlapping vibrations of unordered organic matter in thermally matured fossils (*26-29, 32-33, 37*). Carbonaceous fossils do not contain graphite, as demonstrated by mass spectrometry and light spectroscopic studies of fossil organic matter (see Tab. 1 and references therein). Broad D- and G-bands in spectra of carbonaceous fossils are associated with -C-C-, -C=C- (and -C-N-) functional groups (*3, 26-33, 37*). These are well established signatures (*3-15, 26-33, 37*) that co-occur with those of other major functional groups in Raman spectra of heterogeneous carbonaceous tissues (*3-15*). Nevertheless, the -C-C-/-C=C-(around 1350 cm^-1^) and -C=C-/-C-N-bands (1600 cm^-1^) associated with unordered carbonaceous materials (*3-15, 26-33, 37*) were interpreted as artefacts by Alleon et al. (*21*) in the two new fossil spectra they analyzed (*21*, fig. 3f). In contrast these bands were considered significant in the Raman analysis of a carbonaceous fossil (*31*, fig. 9c) involving one of the same authors.

**Table 1:**
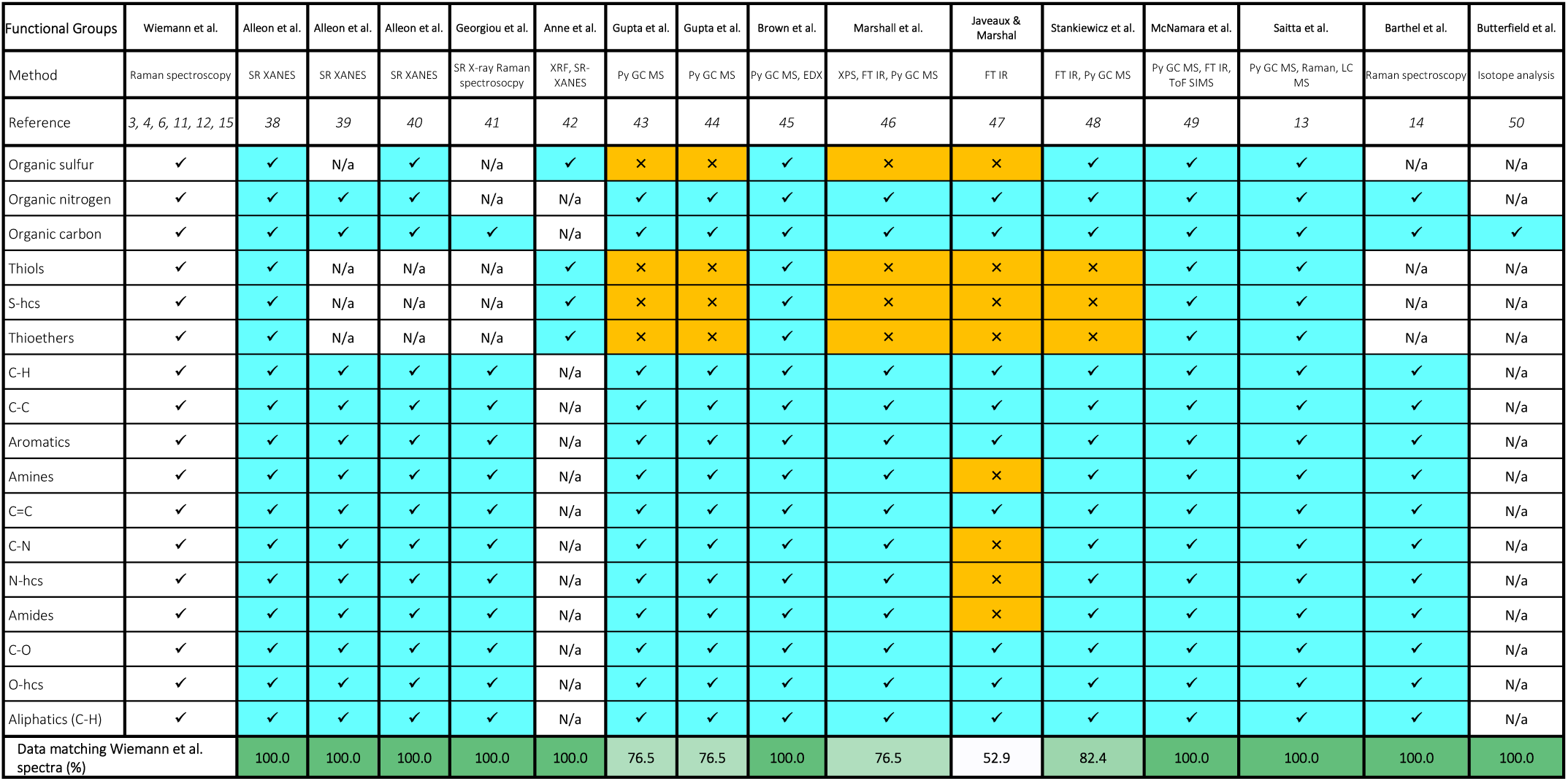
Assessment of a diversity of paleontological and geochemical studies on the composition of fossil organic matter. Details of the authors of each study, the method used, and the correspondence to functional groups identified in (*3, 4, 6, 11, 12, 15*), as well as all references to these studies are provided. The positive identification of the listed functional groups is indicated by blue cell colors, while the absence of a given functional group (in case of the studies by Gupta et al. this is a result of the methodology) is indicated by an orange cell color. Not all studies assessed the complete suite of functional groups listed here, as indicated by a white cell color and the statement ‘N/a’. The last row of the table calculates how well the assembled functional group data for each study match the disputed studies, and cell colors follow a gradient from white to green depending how well the results of each independent study matches data in the disputed studies (*3, 4, 6, 11, 12, 15*).

The spectra that Alleon et al. (*21*) regarded as invalid include not only D- and G-bands for metamorphosed fossils, such as Burgess Shale specimens (*11*), and -C-C-/-C=C- and -C=C-/-C-N-bands in less pressure- and temperature-affected (P/T-affected) specimens (*21*, fig. 1, dark red areas) but also sharp crystal lattice vibrations associated with minerals (*21*, fig. 1, calcite peak - compare to *25*) confirming that decomposition and reconstruction of baselined spectral data based on wavelet transformation provides little information on noise contribution (*22-24*). Nonetheless, the wavelet transform analysis and maps of a few spectra are the sole evidence presented for rejecting all Raman data in six independent studies (*3, 4, 6, 11, 12, 15*) and, by implication, the application of Raman spectroscopy to organic materials in general.

### 3.3. 3-D ChemoSpace analyses reveal no similarity between edge filter artefacts *(21*) and the disputed Raman data

Alleon et al. (*21*) compared their own Raman spectra (n=5) to those of the Mazon Creek shrimp *Acanthotelson* (*12*) and the eggshell of a modern *Rhea* (*4*) (*21*, fig. 1) by means of frequency decomposition alone. Their five raw spectra (*21*, figs. 3d, e) show broad bands without evidence for sharp mineral or organic peaks, and were baselined. Alleon et al. (*21*) based their assertion that our published Raman spectra are invalid largely on superficial similarities in the contribution of 64 cm^-1^ and 128 cm^-1^ frequency components to their baselined *Peachocaris* spectrum, the ‘Semrock’ edge filter spectrum, and our two spectra (see Section 3.1-3.2 on issues with this approach). They argued, based on their frequency contribution maps, that the disputed spectra resemble only edge filter ripples (see Section 3.1-3.2) and claimed, based on similarities with their own, processed data (*21*, figs. 3d, e) that the adaptive baseline used for the disputed spectra only exaggerates edge filter artefacts. However, they offered no quantitative comparisons between their own raw and baselined spectra and the disputed ones, nor did they plot the spectra together for comparison.

In order to provide a quantitative assessment, we compiled all the raw spectra (n=5) and their adaptively baselined equivalents (n=5) based on source data provided by Alleon et al. (*21*, data on Dryad), together with our raw data on Mazon Creek fossils (n=31, *12*), new spectra of Mazon Creek sediments (n=9), and eggshells (n=18, 13 of which are fossils, *4, 15*). We baselined (adaptive, 30%) and normalized all these raw data together, selected relative intensities at n=26 band positions (24 organic bands published in *11* together with phosphate – 954 cm^-1^ and carbonate – 1085 cm^-1^, see *24*), and compiled the data into taxon character matrices. We analyzed our Mazon Creek fossil, sediment and eggshell spectra against: 1. the raw spectra presented by Alleon et al. (n=5) (Figs. 4a, d, g); 2. their raw and baselined spectra (n=5+5) (Figs. 4b, e, h); and 3. their *Peachocaris* spectrum (Figs. 4c, f). We compared our spectra to their five raw spectra to assess the degree of compositional overlap (Figs. 4a, d, g). We also tested whether processing their already baselined spectra significantly alters the position of a sample in the 3-D ChemoSpace in order to assess the influence of (excessive) adaptive baselining (not tested for in *21*) on ChemoSpace clustering (Figs. 4b, e, h). An additional 3-D ChemoSpace Principal Component Analysis (PCA) includes only their Mazon Creek *Peachocaris* – the only spectrum they subjected to frequency contribution mapping – in order to compare our spectra to the one that they analysed and dismissed as artefactual (Figs. 4c, f).

Each character matrix was subjected to a Principal Component ChemoSpace Analysis in PAST 3.0, and the results were exported (Fig. 4). In addition to this, all our data were separately combined with the raw spectra presented by Alleon et al. (n=5) (Fig. 4i), and with their raw and baselined spectra (n=10) (Fig. 4j), and subjected to Discriminant Analyses differentiating between their spectra and ours. None of the 3-D ChemoSpace or Discriminant Analysis plots (Figs. 4a-j) show overlap between spectra considered by Alleon et al. (*21*) to between spectra considered by Alleon et al. (*21*) to contain artefacts and ours (*4, 12, 15*). Regardless of the degree of adaptive baselining applied, their data do not overlap with ours (Figs. 4b, e, h), countering their assertion that adaptive baselining affects the conclusions drawn in the disputed, quantitative studies (*4, 12, 15*). Notably, the data presented by Alleon et al. (*21*) do not resemble the biological or mineralogical separation of samples observed in the disputed data (*4, 12, 15*), but are spread across the ChemoSpace (Fig. 4i).

### 3.4. Raman spectra match those from other laboratories and show substantial compositional variability

We plotted spectra from the publications disputed by Alleon et al. (*21*) and compared them to spectra independently obtained by a different laboratory (*14*) and to the 64 cm^-1^ and 128 cm^1^ edge filter spectrum frequency components presented by Alleon et al. (*21*, figs. 1,2). Spectra obtained under comparable analytical conditions for different samples differ, as evidenced by the Raman peak and band representation (Fig. 5). Compositional information is encoded along the x-axis of the Raman spectral plot (Fig. 5), and peaks or bands at a given Raman shift characterize a specific functional group or crystal lattice vibration. Generally, the use of different instruments, excitation laser wavelengths, and the surface properties of samples may impact the composition of the resulting spectra.

**Fig 5:**
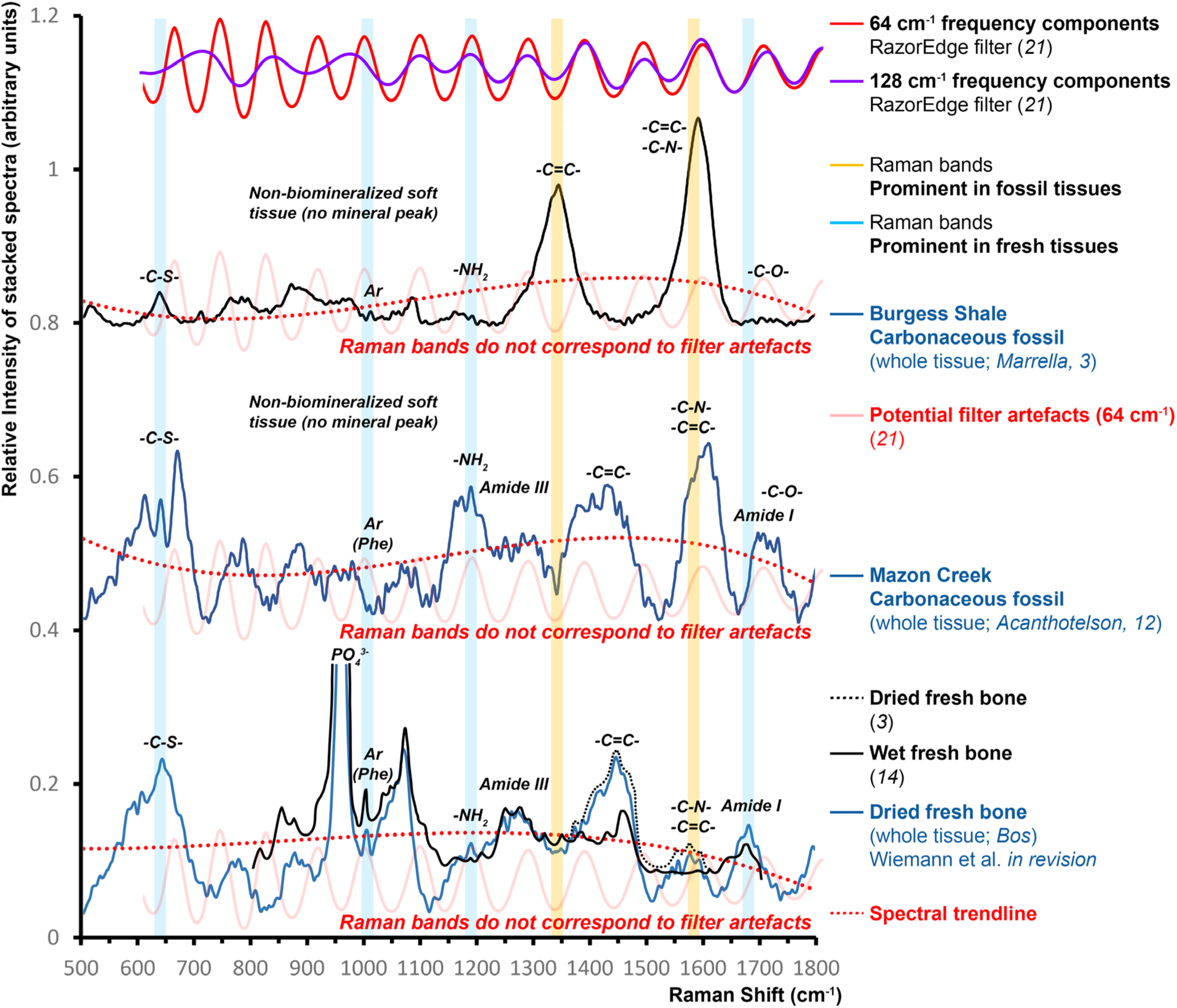
Raman spectra do not resemble filter artefacts. The fossil spectra were obtained with 10 technical replicates and subjected to adaptive baseline subtractions. Raman bands that are prominent in fresh tissues (blue bands) such as *Bos* bone decrease in relative intensity with diagenetic modification of organic matter, whereas bands associated with -C=C- and -C-N-bonds (yellow bands) increase in relative intensity in fossils.

The spectrum we obtained for dried modern bison bone (Fig. 5) matches all major peaks present in the independently published bone whole-tissue spectrum (from *14*; see also *5, 7-10, 13-14, 26-33, 37*). Even weak organic signals, such as that at 1000 cm^-1^ ± 2 cm^-1^ [*Ar* (*Phe*) in Fig. 5] associated with aromatic amino acid residues in modern bone proteins (primarily phenylalanine), are clearly represented, reflecting the high quality of the spectral data obtained (*14, 35, 36*). There are no published whole-range *in situ* Raman spectra of carbonaceous Mazon Creek fossils beyond those in McCoy et al. (*12*), so no independent comparison is available. However, our spectrum of *Marrella* from the Burgess Shale (Fig. 5) resembles published spectra of pressure- and temperature-metamorphosed carbonaceous fossils from the Burgess Shale and elsewhere (*26-33, 36*). The three spectra in Fig. 5 show differences characteristic of progressive diagenetic alteration (compare *29*), resulting in the loss of organic nitrogen, oxygen-, and sulfur-bearing groups in favor of a more ordered organic carbon scaffold in the Burgess Shale fossil.

Neither the spectra in Fig. 5, nor any of the others (*3, 4, 6, 11, 12, 15*) claimed as artefactual (*21*), show evidence of the sinusoidal pattern characteristic for edge filter ripples.

### 3.5. Raman peak evolution during experimental oxidative maturation mirrors fossilization

Alleon et al. (*21*) overlooked our work on tracing the transformation of Raman peaks/bands through experimental maturation (e.g., *3*, fig. 2). Band shifts reflect the effect of alteration and validate signals in the disputed analyses by revealing their origins (*3, 4, 6, 11, 12, 15*). This is illustrated by published data for untreated fresh, decalcified, experimentally oxidized, and fossil eggshells. *Rhea* eggshell has a thick organic scaffold characteristic of all paleognath eggs with barely detectable trace amounts of pigment (*4*). Raman band shifts resulting from oxidation experiments on eggshell organics (*3*) show a similar trend to the composition of organics in fossil compared to modern eggshells (Fig. 6). Acid-based eggshell decalcification leads to the loss of biomineral crystal lattice vibrations (CO_3_ ^2-^ band at 1085 cm^-1^ Raman shift) in decalcified spectra, but has only minor effects on spectra of the organic scaffold (Fig. 6). Experimental oxidation of extracted eggshell organic scaffolds leads to a relative increase in -C=C- and -C-N-functional groups, coinciding with the loss of amine-bearing amino acid residues (Fig. 6; see *3*, fig. 4b, c for details). This alteration occurs in various tissue types with only minor differences (*3*), and has been reproduced for larger, independent sample sets (*11*). Net enrichment plots that subtract spectral signatures of modern tissues from their fossil homologues reveal that fossilization triggers a relative increase in the abundance of unsaturated carbon bonds, N-, O-, S-heterocycles, ethers, esters, and thioethers, while causing a net depletion in nucleophilic amino acid residues (*3, 11*). Comparable patterns of compositional change during fossilization have been demonstrated not only in vertebrate eggshells, but also in bones, teeth and enamel scales, invertebrate soft tissues, and invertebrate biomineralized tissues (*3, 4, 11*) and the transformation of spectral peaks/bands during fossilization has been corroborated through ChemoSpace PCAs of spectra of unaltered modern, experimentally oxidized, and fossil tissues (*3, 4, 11*, but also *33*). Fossilization experiments produce signatures similar to those of fossils, and generate intermediates in a ChemoSpace PCA which overlap both modern and fossil clusters (*3*, fig. 4b). These spectra do not show evidence for periodic bands corresponding in shape or spacing to sinusoidal edge filter artefacts (Fig. 6). Bands based on the same filter artefacts (*21*) would have an identical band representation, and would not show patterns consistent with diagenesis identified by spectroscopy (Figs. 5, 6) and statistical analyses (*3, 4, 6, 11, 12, 15*), and independent assessments (Tab. 1).

**Fig 6:**
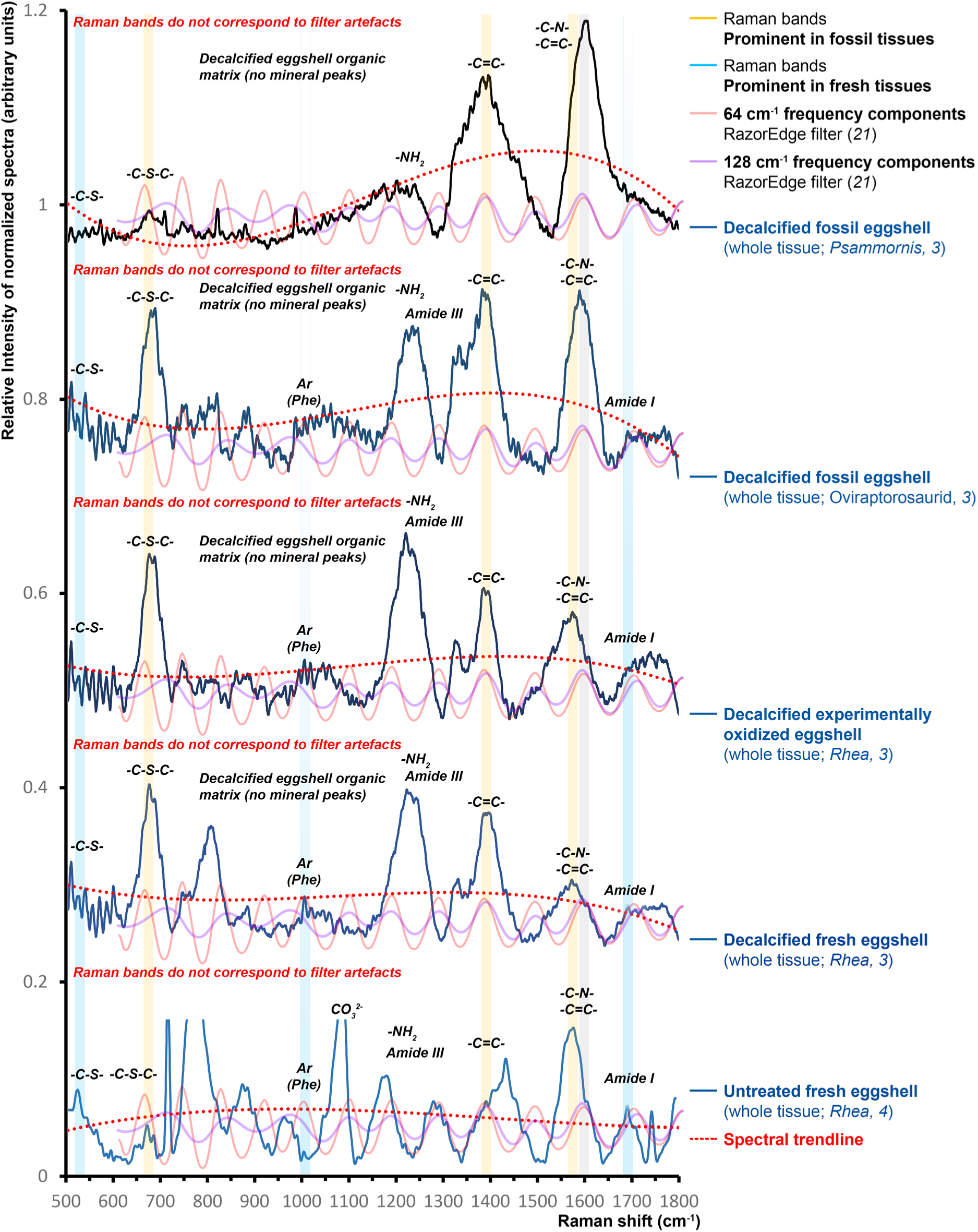
Raman spectra of modern, experimentally oxidized and fossil eggshell organic matter (*3, 4, 11*) show peak/band evolution throughout progressive tissue transformation. All spectra were acquired for whole-tissues with 10 technical replicates, normalized, and subjected to an adaptive baseline subtraction (30%). Changes in spectra from untreated, fresh (bottom) to fossil (top) eggshell, illustrate how certain organic bands (which are generally broad due to the highly heterogenous composition of whole-tissue samples) increase at different stages of maturation, while others systematically decrease. The compositional transformation of modern *Rhea* eggshell organics to fossil *Psammornis* eggshell organic matter can be traced band by band.

### 3.6. The disputed Raman spectra cluster in statistical analyses in chemically, biologically and geologically meaningful ways

The assertion (*21*) that published spectra are artefactual, predicts that they do not contain chemically, geologically, or biologically meaningful information. In that case multivariate statistical analyses (*17-20, 24*) would not yield consistently and reproducibly meaningful clustering of samples based on their spectral composition (*3, 4, 6, 11, 12, 15*). However, quantitative analyses of sets of our published Raman data show that they reproducibly separate into 1. sedimentary vs. fossil organic matter (*3, 4, 6, 11, 12, 15*), 2. modern samples vs. experimentally matured vs. fossil samples (*3, 11*), 3. biomineralized vs. non-biomineralized samples (*11, 15*), 4. different tissue types (*11, 12, 15*), 5. animals with different metabolic capabilities (*11*), 6. clades (*11*), and 7. pressure and temperature conditions experienced during diagenesis (*11, 26-28, 32, 33*). Biological signals retained in the molecular composition of fossils always resemble signals in corresponding extant taxa (*3-6, 9, 11, 12, 15*). Some of these molecular signatures have also been detected in independent studies using different analytical methods (e.g., the tissue type signal reproduced by Alleon et al. *30*, or thermal maturation signals in *26-29, 32, 33*).

All Raman spectra include spectral noise which commonly affects the shape of the baseline (see *33* for fossil examples). The application of a common adaptive baseline to different kinds of analyzed samples might affect some normalized spectra more than others: However, any impact this might have on interpretations is circumvented by evaluating large spectral data sets statistically (as in *3, 4, 6, 10-12, 15*) using ChemoSpace Principal Component or Discriminant Analyses (*16, 17, 20, 24*). The conclusions of the papers (*3, 4, 6, 11, 12, 15*) that Alleon et al. (*21*) disputed were the result of such ordination methods applied to spectral intensities obtained at previously identified, informative band positions. The routine used in the disputed papers corresponds to the multivariate analysis of Raman spectra in whole-tissue cancer diagnostics (*16, 18-20*), and the investigation of whole-tissue archeological remains (*33*). Such a statistical approach is a more robust route to generalized inferences than relying on qualitative analyses of small numbers of samples (*16-20*).

Even in a hypothetical scenario where bands in identically treated spectra included a certain contribution from fluorescence, resulting in broadened spectral shoulders, the targeted analysis of informative bands would prevent any such noise from impacting a comparative ChemoSpace analysis (*16, 17, 20, 24*). Similar analyses are commonly applied to data obtained using instrument parameters comparable to those that yielded the results that Alleon et al. (*21*) considered invalid (*16-19, 32*). Those too focus on selected relative intensities to eliminate the impact of so-called ‘horse-shoe effects’ resulting from the analysis of whole-spectral data containing potentially broad organic bands (*17-19, 32*).

### 3.7. Raman maps are consistent with the nature of biological and geological materials

Raman maps published in the disputed studies (*3, 4, 6, 11, 12, 15*), which reflect the compositional data revealed by spectra, were overlooked by Alleon et al. (*21*). Individual bands mapped onto fossils revealed signals that are spatially constrained to fossil organic matter in extracted, fossil extracellular matrix (ECM) patches (*3*, fig. 3), *in situ* maps of modern avian, dinosaur and stem bird protoporphyrin egg spots (*4*, fig. 2 and supplementary information), and *Mussaurus* eggshell embedded in sediment (*15*, supplementary fig. 2).

Raman maps of ‘artefactual bands’ would not be expected to follow the outline of extracted ECM patches (*3*, fig. 3a) if spectral signatures were just composed of edge filter ripples as suggested by Alleon et al. (*21*). Nor would they abruptly stop at the eggshell-sediment interface (*15*, supplementary fig. 2) if there were no difference in signal between carbonaceous material and sediment as suggested by indistinct artefactual spectra of fossils and sediments in Alleon et al. (maps in *15*, supplementary fig. 2; compare to figs. 3d,e in *21* and our ChemoSpace in Fig. 4i). Differences in the spatial distribution of N-heterocycles and less abundant amide I functional groups (*3*), or thioethers (*15*, supplementary fig. 2) require an original chemical signature. Raman signals that follow biological structures and show variation in the distribution of different functional groups cannot be artefactual.

### 3.8. The results of Raman analyses match those of independent cross-methodological assessments

Alleon et al. (*21*) further claimed that our spectra of fossil carbonaceous materials can only be artefactual, as fossil organic matter cannot contain functional groups beyond -C-C- and -C=C-units associated with graphite-like D- and G-bands in unordered organic materials (*21*), significant signals that, unexpectedly, were also identified as noise by their wavelet transform analysis.

The assumption that fossil organic matter cannot be more complex in composition is at odds with the published literature: it is well known that fossil organic matter is characterized by a heterogenous composition which converges in time due to pressure-, temperature-and redox-induced alterations (*3, 11, 26-29, 32, 33*). Fresh tissues yield a compositional diversity represented by numerous organic bands (*3, 11, 34, 36*). Depending on the diagenetic history, fossil organic matter is more or less crosslinked as a result of oxidative alteration (*3, 11, 26-29, 32, 33*). Such differences in composition are mirrored in the different appearance and properties of fossil soft tissues: Late Paleozoic and Mesozoic fossil soft tissues (e.g., *3*, fig. 1) differ from highly matured silvery-opaque carbonaceous films that are characteristically simpler in chemical composition, by exhibiting a transparent-to-opaque brown color and pliable rheology (*3-15*), resembling their source tissues morphologically. Such less altered, 3-D fossil soft tissues have undergone little maturation and are usually more complex in composition.

Wiemann et al. (*3, 11*) demonstrated statistically that biomolecule fossilization is characterized by the loss of nucleophilic amino acids residues and fatty acid and sugar moieties, reaction educts that are consumed during the formation of N-, O-, S-heterocycles. Fossil organic matter subjected to pressure- and temperature maturation is further characterized by the progressive loss of N-, O-, S-containing functional groups in favor of planar, aromatic structures (*3, 11, 26-29, 32, 33*). Numerous studies (*13, 14, 38-50*) report the composition of a diversity of fossil organic matter obtained from samples covering a taxonomic range, age and diagenetic history comparable to samples analyzed in the disputed studies (Tab. 1). These independent, cross-methodological analyses identify a similar diversity of functional groups to that inferred from the Raman spectra of the disputed studies.

The different functional groups detected through Raman spectroscopy (*3, 4, 6, 11, 12, 15*) and regarded as invalid by Alleon et al. (*21*) have been reported in numerous independent studies using Raman or Infrared spectroscopy (FT IR), Pyrolysis Gas Chromatography mass spectrometry (Py GC MS), and Synchrotron radiation X-ray absorption near edge structure spectromicrocroscopy (SR XANES). Based on the requirement for targeted organic phase extractions, Py GC MS studies often contain little information of organo-sulfur compounds in fossil organic matter (see *42* and *43*, in Tab. 1) but this may not reflect their original absence. Some of the best matches to the composition of fossil organic matter reported in the disputed studies (*3, 4, 6, 11, 12, 15*) are, surprisingly, presented in papers by Alleon et al. (*37-39*). For example, using Synchrotron Radiation XANES, Alleon et al. (*30*) detected all the functional groups identified through our statistical analyses of the disputed Raman spectral data.

## 4. Discussion and conclusions

Alleon et al. (*21*) used an unconventional analysis of two out of more than 200 published spectra of modern animal tissues, carbonaceous fossils and associated sediments (*3, 4, 6, 11, 12, 15*) to argue that Raman spectroscopy cannot be used to identify patterns in the composition of fossil organic matter. Their conclusion relies primarily on superficial similarities in maps of spectral frequency contributions, and the resemblance of decomposed and summed spectra to their original source spectra.

We compared frequency contribution maps (Fig. 1) and spectral plots from the disputed studies (Figs. 2, 3, 5, 6) with the edge filter ripples highlighted by Alleon et al. (*21*). We highlighted the diversity of spectra included in the studies they dispute, and showed how experimental studies explain the evolution of Raman peaks during fossilization, all aspects that they (*21*) did not address. Based on the resulting insights, we discussed how the wavelet transform analysis used by Alleon et al. (21) is unsuitable for identifying spectral noise. The sum of any two substantially distinct spectral frequency components with a wavelength shorter than the base width of the broadest Raman band will closely resemble the original spectrum. Subtracting such a summed function from the original source spectrum will, depending on the frequencies chosen, eliminate even sharp mineralogical peaks (see *21*, fig. 1a). Indeed, Alleon et al. (*21*) identified sharp and intense mineral crystal lattice vibrations (CO_3_ ^2-^ peak of calcite in the *Rhea* spectrum), as well as established D-/G-bands (in P/T-matured fossils) and -C-C-/-C=C- and -C=C-/-C-N-vibrations (in less matured samples; *3, 11, 26-29, 32, 33, 37*) as noise – an anomaly that demonstrates the inappropriate nature of their approach. The interpretations of Alleon and colleagues are not consistent: while broad D- and G-bands in the disputed spectra (Figs. 5, 6; *3, 4, 6, 11, 12, 15*) were considered artefactual (*21*), the same bands are considered significant and informative in their own papers (*30, 31*) where no corrections for potential edge filter ripples were applied (*21*).

The conclusions of Alleon et al. (*21*) rely exclusively on qualitative comparisons; they offered no identically scaled (i.e., directly comparable), or quantitative comparisons between the edge filter spectra, the two disputed spectra, their new analyses, or their frequency contribution patterns. In order to remedy this omission (Figs. 1, 2) we presented the frequency contribution maps at the same scale (*21*, figs. 1a, c, 2b). Direct comparisons refute the suggestion (*21*) that these plots show similarities. Likewise the 64 cm^-1^ and 128 cm^-1^ frequency components calculated from our two spectra (*21*) do not resemble each other (Fig. 2). We baselined and normalized the spectra presented by Alleon et al. (*21*) which, combined with data from the disputed studies, generated a data matrix of n=68 spectra (sourced from *4, 12, 15*). Our plots (Figs. 3, 5, 6) demonstrate that the spectra presented by Alleon et al. (*21*) show no resemblance to our data – the artfactual, smooth edge filter ripples are offset from the bands in the fossil spectra (Fig. 3). None of the sharp organic and mineral peaks combined into broad organic bands, which are evident in all our spectra, are present in their spectra (Fig. 3). 3-D ChemoSpace PCA and discriminant analyses (Fig. 4) show no compositional overlap between the disputed spectra of modern, fossil, and sediment samples and the artefactual spectra presented by Alleon et al. (*21*). Using their raw and baselined spectra, together with raw data from the disputed papers, we demonstrate that even exaggerated adaptive baselining (Fig. 5) does not change how samples group in a 3-D ChemoSpace.

Evidence from a range of studies overlooked by Alleon et al. (*21*) provides further validation of the original nature of detected Raman spectral signatures: Raman maps of functional groups in modern and fossil samples reveal that Raman signals are confined to biological morphologies (*3, 4, 15*). ChemoSpace analyses of large spectral sample sets cluster taxa in biologically and geologically meaningful ways that have been reproduced multiple times (*3-6, 9, 10-12, 15, 33*), revealing molecular signatures that resemble those present in modern tissue equivalents. Geochemical and paleontological studies on fossil organic matter composition (Tab. 1, *13, 14, 38-50*) reveal numerous matches between our published Raman data and independent analyses using Raman spectroscopy, X-ray Raman spectroscopy, mass spectrometry, infrared spectroscopy, and Synchrotron XANES studies – including various studies by Alleon and colleagues (*30, 31, 38-40*).

Based on the multiple lines of evidence presented here, we conclude that our spectra contain original, biologically and geologically meaningful molecular information. Statistical analyses of large Raman spectral data sets, as well Raman maps of *in situ* and extracted organic samples, remain powerful tools for the analysis of compositional patterns and their spatial distribution in a variety of geo- and biomaterials.

**Source Data are available from the corresponding author upon request**.

**The authors declare no conflict of interest**.

## Notes

### Competing Interest Statement

The authors have declared no competing interest.

